# Population structure and genetic diversity of chickpea germplasms

**DOI:** 10.1101/2021.08.04.455107

**Authors:** Garima Yadav, Deepanshu Jayaswal, Kuldip Jayaswall, Abhishek Bhandawat, Arvind Nath Singh, Jyotsana Tilgam, Abhishek Kumar Rai, Rachna Chaturvedi, Ashutosh Kumar, Sanjay Kumar

## Abstract

In various leguminous crops, chickpea is the fourth most important legume contributing 3.1% to the total legume production. Grains of chickpea are rich source of proteins, minerals and vitamins which makes them suitable for both food and feed. For any crop to be improved, the knowledge of genetic diversity of wild and elite cultivar is very important. Therefore among various available marker systems, molecular markers are more reliable and accurate, therefore are very commonly used for genetic diversity analysis, phylogenetic studies and cultivar identification. Due to several advantages of Inter Simple Sequence Repeat (ISSR) markers in present study we analyzed the genetic diversity, structure, cross-species transferability and allelic richness in 50 chickpea collection using 23 ISSR markers. The observed parameters such as allele number varied from 3 to 16, and PIC varied from 0.15 to 0.4988 respectively. Further, range of allele size varied from 150 to 1600 bp which shows the significance of ISSR markers for chickpea germplasms characterization. On the basis of ISSR marker genotypic data dendrogram were constructed which divides these 50 chickpea in group I and II showing the reliability of ISSR markers. Among 50 chickpea, the accession P 74-1 is in group I and rest are in group II. Further we made mini-core collection of 15 diverse chickpea and subgrouped them. Dendrogram, PCA, Dissimilarity matrix and Bayesian model based genetic clustering of 50 chickpea germplasms revealed that P 74-1,P 1883, P 1260 very diverse chickpea accession. Characterization of these diverse chickpea would help in maintenance breeding, conservation and in future could be used to develop climate resilient elite cultivar of chickpea. Utilization of these novel ISSRs markers in diversity analysis and population structure characterization of 50 chickpea germplasm suggests their wider efficacy in superior scale for molecular breeding studies in chickpea.

## Introduction

In various leguminous crops, chickpea is the fourth most important legume contributing 3.1% to the total legume production (http://faostat.fao.org). The worldwide production of chickpea was 14.25 million tonnes from 14.7 million ha (http://www.fao.org/faostat/en/#data/QC/visualize). Grains of chickpea are rich source of proteins, minerals and vitamins which makes them suitable for both food and feed. There are total of 43 species has been reported yet under the genus *Cicer* among which only one species i.e., *Cicer arietinum* L. is under cultivation practices and is of economic importance (Sethy *et. al*., 2006a). The genome sequence data revealed the genome size (∼750 Mbp) with 8 basic set of chromosome (Sethy *et. al*., 2006b). International Crops Research Institute for the Semi-Arid Tropics (ICRISAT), Hyderabad, India and International Center for Agricultural Research in Dry Areas, Syrian Arab are major organizations contributing in the collection of chickpea germplasms having 17,258 and 12,647 accessions (Upadhyaya *et. al*., 2008). For any crop to be improved, the knowledge of genetic diversity of wild and elite cultivar is very important. Before 1980s when molecular markers were not known, genetic diversity and relationship were analyzed by biochemical markers. Further, the discovery of molecular markers has made the genetic diversity analysis easy. Among various available marker systems, molecular markers are more reliable and accurate, therefore are very commonly used for genetic diversity analysis, phylogenetic studies and cultivar identification. Due to continuous selection breeding for crop improvement, genetic polymorphisms in the cultivated *Cicer arietinum* L. are very low, hence marker assisted selection (MAS) and marker assisted breeding (MAB) could play very potential role in the development of new varieties. Since past many years, genetic diversity analysis with different molecular markers in various crops has been done efficiently (Ahmad, 1999; Queen *et. al*., 2004; Schulman, 2007; Gill-Langarica *et. al*., 2011; Noormohammadi *et. al*., 2013; Bonman *et. al*., 2015; Mogga *et. al*., 2018; Elshafei *et. al*., 2019; Delfini *et. al*., 2021). Due to several advantages of Inter Simple Sequence Repeat (ISSR) markers over other markers such as PCR based, no requirement of sequence information, distribution across the whole genome, require small quantity of template DNA and cost effective, ISSR markers are extensively used for genetic diversity analysis. The genotypic data of ISSR marker, pedigree analysis and various phenotypic data can assist in the selection of germplasm for further crop improvement through molecular breeding programs (Rao *et. al*., 2007).

In this study we analyzed the genetic diversity, structure, cross-species transferability and allelic richness in 50 chickpea collection using 23 ISSR markers. The observed parameters such as allele number varied from 3 to 16, and PIC varied from 0.15 to 0.4988 respectively. Further, range of allele size varied from 150 to 1600 bp which shows the significance of ISSR markers for chickpea germplasms characterization. On the basis of ISSR marker genotypic data dendrogram were constructed which divides these 50 chickpea in group I and II showing the reliability of ISSR markers. Among 50 chickpea, the accession P 74-1 is in group I and rest are in group II. Further we made mini-core collection of 15 diverse chickpea and sub grouped them. Dendrogram, PCA, Dissimilarity matrix and Bayesian model based genetic clustering of 50 chickpea germplasms revealed that P 74-1,P 1883, P 1260 very diverse chickpea accession. Characterization of these diverse chickpea would help in maintenance breeding, conservation and in future could be used to develop climate resilient elite cultivar of chickpea. The study of genetic diversity and population structure of 50 chickpea and development of 15 chickpea core collection will serve as important knowledge resources for future studies like GWAS, and mapping.

## Material and methods

### Plant Material

Total fifty individuals of wild chickpea germplasm (Supplementary Table 1) were grown in the field of ICAR-IISS, Mau, India. Genomic DNA from freshly procured leaves of chickpea was isolated by CTAB method with minor modification (Murray and Thompson, 1980). The wild chickpea germplasm having varying concentrations of secondary metabolite content and to overcome the inhibitory effect of secondary metabolite content during DNA extraction, the concentration of polyvinyl pyrrolidone K-30 and β-mercaptoethanol were standardized. Further, the quality of extracted DNA was examined over 0.8% agarose gel using lambda uncut marker (Fermentas, Lithuania) and quantified by NanoDrop 2000 (Thermo Scientific, USA).

### Amplification validation and polymorphic potential evaluation

Fifty chickpea germplasm were genotyped by 23 inter simple sequence repeat (ISSR) markers (Supplementary Table- 1) and DNA fingerprint was developed (Table-1). For all the, PCR amplification efficiency with 23 ISSR markers were analyzed in 25µL reaction volume with 20 ng of DNA template. The PCR program initiated by pre-denaturation step at 94°C for 5 min, subsequent 35 cycles of denaturation at 94°C for 1 min, annealing at optimum temperature 50°C for 1 min, primary extension at 72°C for 2 min and the final extension was done at 72°C for 10 min in Thermo Cycler (Eppendorf). The amplified PCR products were separated on 1.5% agarose gel, and amplicon size were estimated based on 100 bp DNA ladder (Genedirex) as a reference.

### Molecular data analysis

The amplification profiles with ISSR markers were scored based on their presence (1) or absence (0) across all individuals (SupplementaryTable-2). The amplified product were categorized as monomorphic, polymorphic and null alleles on the basis of same, different and absence of amplified PCR product across all the individuals. Cross-transferability of the ISSR markers and amplicon size were recorded for all the chickpea germplasms to construct UPGMA dendrogram and Principle Component Analysis (PCA) based on Jaccard’s coefficient using DARwin6 (ver.) software (Perrier and Jacquemoud-Collet 2006). The polymorphic information content (PIC) values were calculated for each marker using online PIC calculator (https://www.liverpool.ac.uk/~kempsj/pic.html) (Table-1). Genetic structures of fifty individuals of chickpea were inferred using Bayesian algorithm-based STRUCTURE software (ver. 2.3.3) (Pritchard et al., 2000; Falush et al., 2007). Further dissimilarity matrix (SupplementaryTable-3) was measured using DARwin6 software (Perrier and Jacquemoud-Collet 2006).

## Result and discussion

Genome of plants has several repeated DNA sequence such as Inter Simple Sequence Repeats (ISSR) and Simple Sequence Repeats (SSRs). In the past, when these sequence repeats were considered as ‘junk’ DNA which have been used for plant genetic diversity analysis but with the time when sequencing technologies evolved, Now it is well known that these sequence repeats are the major part of plant genome regulate gene expression and packing of genomes. ISSRs are usually repetitive DNA sequences mostly 1-6 bases (Litt *et al*., 1989) which are PCR based, do not require prior sequence information, distributed throughout the genome, polymorphic and are cross transferable (Welsh *et al*., 1990, Wang *et al*., 1994). Chickpea (*Cicer arietinum* L.) belongs to the family leguminosae, is the fourth most important pulse crop of the world and India is among the largest producer country (FAO, 2010). Chickpea seeds are nutrient rich due to the presence of balanced protein, carbohydrates, vitamins and minerals with the low levels of anti-nutritional factors (Wang *et. al*., 2010; Santiago *et. al*., 2010). Various biotic and abiotic stresses reduce the chickpea production across the globe, therefore, bringing of desirable traits in elite cultivar of chickpea from crossable donor cultivar is need of the hour (Singh *et. al*., 2008). In this scenario, ISSRs markers could play potential role for comparative genomics study, traits identification, genetic purity, association studies and introgression of desirable traits in chickpea through marker-assisted breeding (Gautam *et. al*., 2016; Jayaswall *et. al*., 2019b; Jayaswall *et. al*., 2019a).

### Scoring and data analysis

Twenty five ISSR markers were selected for identification of polymorphism, cross-species transferability, genetic diversity and population structure analysis in 50 chickpea germplasms. Among twenty five only 23 ISSR markers were amplified at least in one of the individuals of chickpea. The amplification failure in some chickpeas could be due to not binding of primers in the genome. In present study, polymorphism was found to be higher than SSRs due to random binding of ISSR primers to the genome of diverse chickpea germplasms. The observed parameters like allele number varied from 3-16, allele size varied from 150 – 1600 bases and polymorphic information content (PIC) value ranges from 0.15 to 0.4988 (Table 1).

### Understanding of taxon and genetic relationship among 50 chickpea germplasms

In plant molecular breeding, the genetic variability is a primary requirement and further, population structure analysis in the selection of elite diverse germplasms (Chakraborty *et. al*., 2016). The rationale of the genetic structure analysis is to understand the population homogeneousity and genetic variability. Population structure is required for mapping of agronomically important genes and dissection of important traits (Wei et al., 2006).Twenty three ISSRs markers were selected for screening of cross transferability and polymorphism studies in 50 chickpea germplasm. Based on the data from 23 ISSR markers (2 ISSRs were not amplified) dendrogram were constructed that divides 50 chickpea germplasms into two group (group I, & II). In group I, germplasm P 74-1 is highly diverse and remaining 49 chick pea fall in group II which have been further divided in different subgroups. Based on UPGMA dendrogram (Fig.1) we selected diverse 15 chickpea core collection (Table 2). Germplasms P 74-1,P 1883, and P 1260 belongs to I, IIB, IIBb, subgroup respectively(Fig.1). Further in Polymorphic Component analysis (PCA), they fall in cluster I(Fig.2). Additionally in structure analysis they belongs to cluster A (Fig.3). Further it was observed that dissimilarity of P 74-1, P 1883, P 1260 to germplasm P 4051 is 78%, 55%, 54% respectively (Table 2 and Supplementary table 3). Dendrogram, PCA, Dissimilarity matrix and Bayesian model based genetic clustering of 50 chickpea germplasms reveal that P 74-1, P 1883, and P 1260 are very diverse than chickpea germplasms P 4051.

**Fig. 1.**
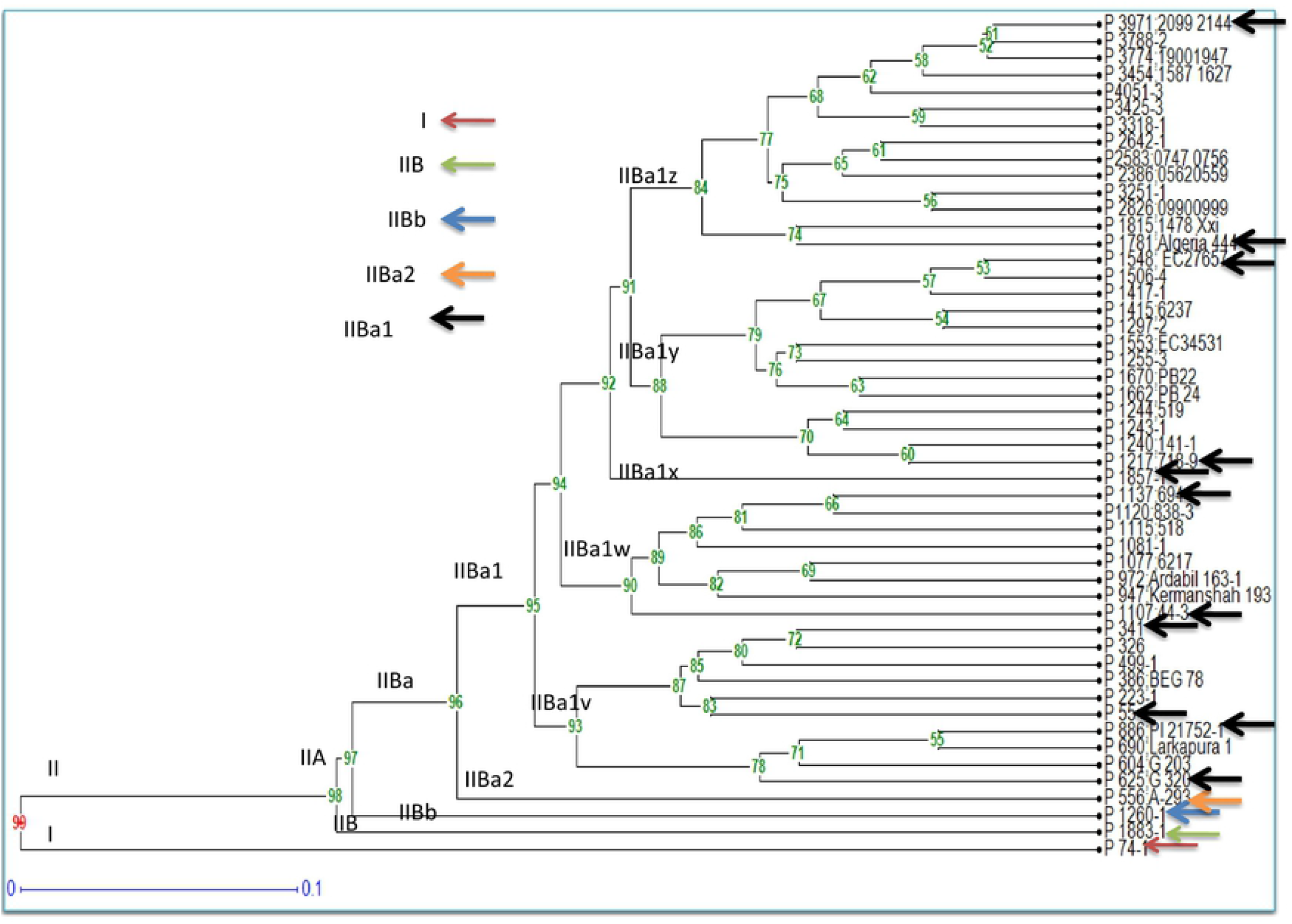
Dendrogram of 50 chickpea germplasms based on 23 inter simple sequence repeats (ISSR) markers

**Fig. 2.**
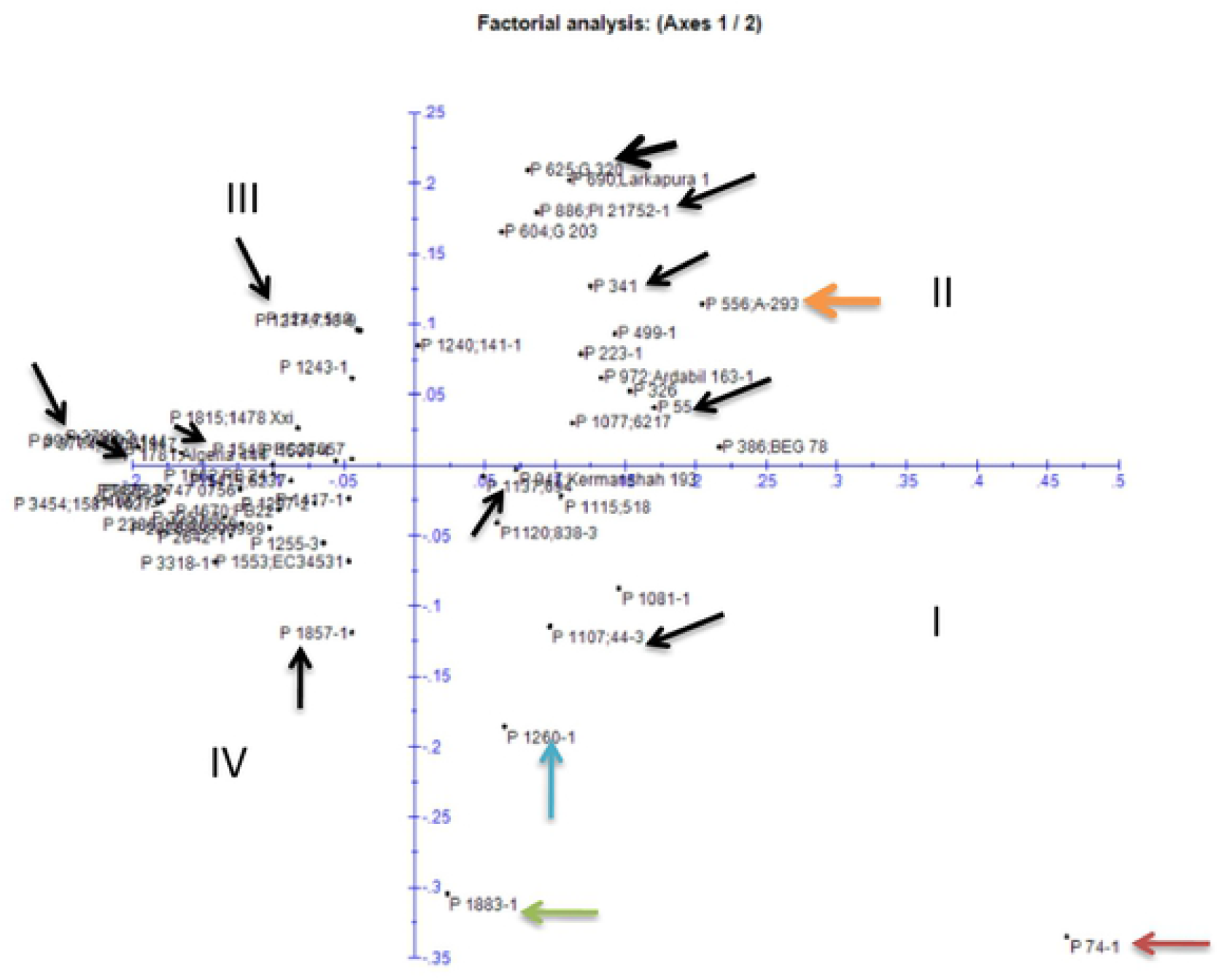
ISSRmarker based principle component analysis (PCA) showing two dimensional distributions of 50 chickpea germplasms

**Fig. 3.**
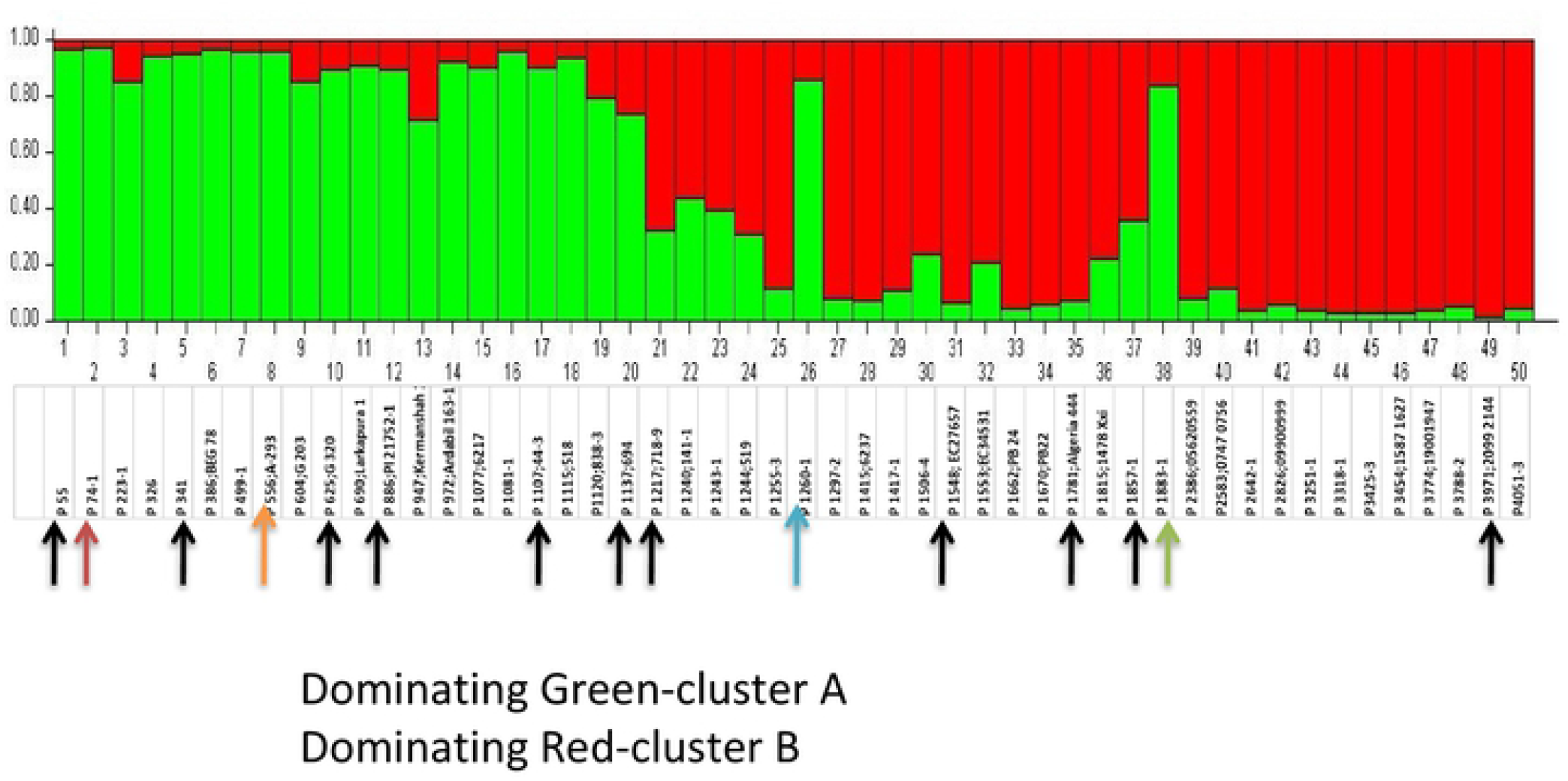
Bayesian model based genetic clustering of 50 chickpea germplasms

Further, P 556, P 625, P 886, P 55, P 341 belongs to IIBa2, IBa1v, IBa1v, IBa1v, and IBa1v group respectively. Germplasm P 556, P 625, P 886, P 55, and P 341 falls in cluster II(Fig.1). P 556, P 625, P 886, P 55, P 341 germplasm also belong to cluster A of Bayesian model based genetic clustering of 50 chickpea germplasms(Fig.2). Additionally P 556, P 625, P 886, P 55, P 341 has dissimilarity 46%, 40%, 40%, 40%, and 40%, respectively(Table 2 and Supplementary table 3) from chickpea germplasm P 4051. P 1107 and P 1137 which belongs to cluster IIBa1w of UPGMA dendrogram(Fig.1).Additionally P 1107 and P 1137 belong to group I of two dimensional distributions of 50 chickpea germplasms(Fig.2).Germplasms P 1107 and P 1137 belong to cluster A of Bayesian model based genetic clustering of 50 chickpea germplasms(Fig.3). P 1107 and P 1137 both have dissimilarity 39% from chickpea germplasm P 4051(Table 2 and Supplementary table 3).P 1857 falls in group IIBa1x of UPGMA dendrogram (Fig.1).and belongs to IV cluster of two dimensional distributions of 50 chickpea germplasms(Fig.2). It belongs to cluster B of Bayesian model based genetic clustering of 50 chickpea germplasms(Fig.3).Its dissimilarity is 35% from chickpea germplasm P 4051(Table 2 and Supplementary table 3). Germplasm P 1217, P 1548, P 1781, P 3971 belongs to group IIBa1w of UPGMA dendrogram(Fig.1).Germplasms P 1217, P 1548, P 1781, and P 3971 belongs to cluster III of two dimensional distributions of 50 chickpea germplasms(Fig.2).P 1217, P 1548, P 1781, and P 3971 germplasms fall in cluster B of Bayesian model based genetic clustering of 50 chickpea germplasms(Fig.3).All P 1217, P 1548, P 1781, P 3971 has dissimilarity 33%, 33%, 28% and 16% respectively from chickpea germplasm P 4051(Table 2 and Supplementary table 3). So dendrogram of 50 chickpea germplasms based on 23 inter simple sequence repeats (ISSR) markers (Fig.1), ISSR marker based principle component analysis (PCA) showing two dimensional distributions of 50 chickpea germplasms (Fig.2), Bayesian model based genetic clustering of 50 chickpea germplasms (Fig.3), triangle plot of 50 chickpea germplasms (Fig.4) reveals that all 15 core collection of chickpea are very diverse and could be used to molecular breeding by utilizing these ISSR markers.

**Fig. 4.**
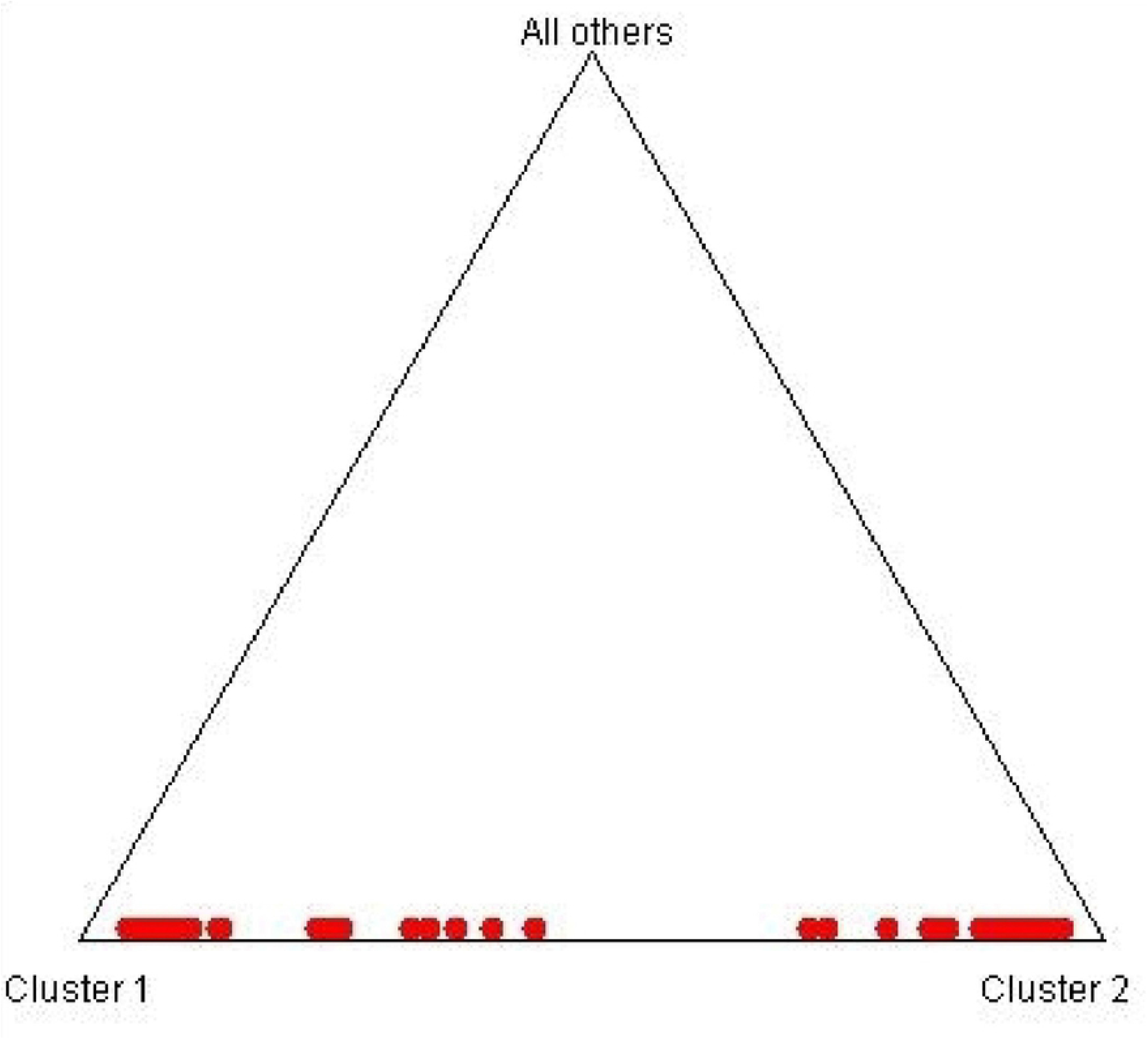
Trianle plot of 50 chickpea germplasms

Therefore information about relatedness of various chickpea within and between group I, and II provides radiant prospect to bring various agronomically desirable traits such as climate resilience genes from donor chick pea to elite cultivated chickpea through marker assisted selection and breeding. Characterization of these diverse chickpeas would help in maintenance breeding, conservation and could be used in future to develop climate resilient chickpea to assist in food security.

## Conclusion

ISSRs in genome influence activity and function of other nuclear and organelle coding gene due to their repeat length. Unfortunately, ISSR marker resources for chickpea have not been well harnessed for genotypic improvement of chickpea. Therefore, a set of 23 ISSRs have been used to expedite molecular breeding of chickpea. High cross transferability and polymorphism of these ISSR markers further reveal their novelty. Utilization of these novel ISSRs markers in diversity analysis and population structure characterization of 50 chickpea germplasm suggests their wider efficacy in superior scale for molecular breeding studies in chickpea. The study of genetic diversity and population structure of 50 chickpea and development of 15 chickpea core collection will serve as important knowledge resources for future studies like GWAS, and mapping.

## Table Legends

Table 1**-**List of primers and their amplification characteristics

Table 2-Panel of core collection of 15 chickpea germplasms

## Supplementary Table Legends

Supplementary Table 1- Detail of samples used for validation DNA fingerprinting of 50 chickpea germplasms

Supplementary Table 2- DNA fingerprint of 50 chickpea germplasms with 23 ISSR markers

Supplementary Table 3- Dissimilarity matrix based on Jaccard’s similarity coefficient for the studied chickpea genotypes

## Acknowledgements

Financial assistance for research by Indian Council of Agricultural Research-Indian Institute of Seed Sciences, Mau is gratefully acknowledged. Further ICRISAT, Hyderabad is gratefully acknowledged for providing chickpea germplasms.

## Compliance with ethical standards

**NA**

## Conflict of interest

The authors declare that they have no conflict of interest.

## Notes

### Competing Interest Statement

The authors have declared no competing interest.

